# Inhibition of acute lung inflammation by a neuroimmune circuit induced by vagal nerve stimulation

**DOI:** 10.1101/2025.02.12.637880

**Authors:** Kaitlin Murray, Michael Cremin, Emmy Tay, Kristina Sanchez, Sierra Schreiber, Ingrid Brust-Mascher, Wesley Leigh, Jannat Ashfaq, Melanie G. Gareau, Colin Reardon

## Abstract

Vagus nerve stimulation (VNS) has been shown to limit immune cell activity across several pathologies ranging from sepsis to auto-immune diseases. While stimulation of vagal efferent neurons has been previously demonstrated to reduce maladaptive host responses during endotoxemia, only selective stimulation of vagal afferent neurons was able to inhibit TLR7-induced macrophage activation and neutrophil recruitment in the lung. These anti-inflammatory actions are facilitated by systemic increases in epinephrine, as VNS significantly increased epinephrine in the serum and bronchoalveolar lavage fluid, and inhibition of epinephrine production eliminated the protection afforded by VNS. Selective afferent VNS induced activation in the nucleus tractus solitarius and the rostral ventrolateral medulla. Inhibition of neuronal activity in this brain region that controls peripheral sympathetic nervous system activity rendered VNS ineffective. Activation of the β_2_-adrenergic receptor (β_2_AR) is critical for innate immune cell suppression, as the anti-inflammatory effects of VNS were eliminated in β_2_AR-knock out mice, and with pharmacological inhibition of the β_2_AR. Analysis of the immune cells responding to R848 critically identified that plasmacytoid dendritic cells were refractive to inhibition by VNS, and this corresponded to lack of β_2_AR expression. These findings demonstrate a novel neuro-immune circuit elicited by VNS that can control acute lung inflammation.

**Summary:** We have identified a novel neuro-immune circuit activated by afferent vagus nerve stimulation to reduce acute lung inflammation. This effect was dependent on vagal-induced adrenal gland-derived epinephrine release that initiates anti-inflammatory β2-adrenergic receptor signaling in innate immune cells within the lung.

## Introduction

Host maladaptive lung inflammation can be caused by a variety of insults ranging from infection with bacterial or viral pathogens to mechanical ventilation. Although coordinated innate immune responses are a critical component of host protection, overactive and aberrant responses can cause significant morbidity and mortality due to lung injury, organ dysfunction, and death^1–3^.

The nervous system is increasingly recognized to actively interface with the immune system to control inflammation in the mesenteric lymph nodes, spleen, intestine, skin, and lungs^4–12^. The lung is innervated by sympathetic, parasympathetic, and sensory neurons that control not only key aspects of physiology but also aid in immune coordination. Lung innervation arising from the vagus nerve has been identified to reduce lung inflammation^13^; however, the precise manner through which immune cell activation is controlled, and which neurons are required, has not been clearly identified. Protective effects of vagal nerve signaling have often been ascribed to the cholinergic anti-inflammatory pathway (CAIP). As a reflex, the CAIP detects systemic inflammation by vagal afferent (sensory) fibers resulting in the activation of vagal efferent (motor) fibers and consequently, sympathetic neurons that project into the spleen and mesenteric lymph nodes (MLN)^14^. Norepinephrine (NE) released from these neurons induces acetylcholine (ACh) release from unique T-cells expressing the enzyme choline acetyltransferase (ChAT) ^4,15,16^. T-cell-derived ACh activates nicotinic alpha 7 receptors (α7R) on macrophages, blocking NF-κB signaling and transcription of pro-inflammatory cytokines such as TNFα^17–19^. This neuro-immune regulatory circuit can also be activated by electrical or optogenetic vagal nerve stimulation (VNS) to reduce LPS-induced systemic inflammation^4,15^. We have demonstrated there is at least one other discrete neuroimmune circuit in the spleen and MLN in addition to the CAIP^16^.

In the context of lung inflammation, while vagotomy increases inflammation during bacterial pneumonia, it is unclear if this effect is due to the loss of the CAIP or the loss of vagal afferent signaling^20,21^. Similarly, experiments using α7R agonists to reduce inflammation during bacterial pneumonia^20^ or ventilator-induced lung injury and inflammation^22^ are not direct evidence of the CAIP but are instead evidence of the direct effects of these compounds. Electrical stimulation of vagal efferent neurons to induce CAIP reduced LPS-induced TNF production in the spleen but not the lung^23^. Although the role of the CAIP in lung inflammation is not clear^24^, electrical stimulation of the intact vagus significantly reduces immune cell recruitment and proinflammatory cytokine production during ventilator-induced mechanical injury^25^. Despite these findings, the precise neuroimmune circuit and mechanisms of action that control acute lung inflammation have not been previously elucidated.

Here we demonstrate that acute TLR3 or TLR7 agonist-induced lung inflammation by instillation of Polyinosinic-polycytidylic acid (Poly(I:C)) or Resiquimod (R848) respectively is reduced by a unique neuroimmune circuit activated by the stimulation of vagal fibers. This lung anti-inflammatory neuroimmune circuit is distinct from the CAIP, as it is independent of T-cell-derived ACh. Electrical or selective optogenetic vagal afferent neuron stimulation limited acute lung inflammation by preventing proinflammatory cytokine production in alveolar and interstitial macrophages, subsequently reducing immune cell recruitment to the lung. Highlighting a role for the central nervous system in the integration of these signals, vagal afferent activation increased neuronal activation, indicated by cFOS expression, in key brain regions, including the nucleus tractus solitarius (NTS) and the rostral ventrolateral medulla (RVLM). Demonstrating the importance of the RVLM, selective afferent vagal nerve stimulation-induced epinephrine release into serum and into bronchoalveolar lavage fluid (BALF). Control of acute inflammation induced by VNS was abrogated by pharmacological inhibition of epinephrine or surgical removal of the adrenal glands, the predominant source of peripheral epinephrine. Using a combination of expression analysis, selective pharmacological antagonists, and knockout mice, the β_2_AR was identified as a required component of this neuroimmune circuit. Based on these findings, we propose a novel lung anti-inflammatory pathway where stimulation of vagal afferent neurons triggers sympathetic activation and release of epinephrine from the adrenal glands to inhibit TLR3 or TLR7-induced lung inflammation. We further propose that selective neuronal stimulation could be a tool to fine-tune and reduce lung inflammation.

## Results

### Vagus nerve stimulation significantly reduces acute lung inflammation

Intranasal instillation of R848, but not vehicle, induced prolific inflammatory cytokine expression within the lung 1-hour post-challenge **(Fig. S1A)**, and significantly increased BALF protein concentration, indicative of inflammation **(Fig. S1B)**. The vagus nerve was stimulated for 20 minutes, with R848 challenge after the first 10 minutes of stimulation **(Fig. 1A).** Vagus nerve stimulation (VNS) of the right cervical vagus nerve inhibited R848-induced mRNA expression of inflammatory cytokines and chemokines compared to non-stimulated sham controls, including *Tnf*α*, Ccl4, Cxcl1,* and *Ifn*β in the lung **(Fig. 1B)**. Serum TNFα protein induced by R848 was also significantly reduced in mice subjected to VNS **(Fig. 1C).** Flow cytometry conducted on single cell suspensions of the whole lung, identified R848 significantly increased TNFα production in alveolar (CD45+, CD11b-, CD11c+, CD64+, SiglecF+) and interstitial macrophages (CD45+, CD11b+, CD11c+, CD64+, SiglecF+), and increased neutrophil (CD45+, CD11b+, Ly6G+) recruitment **(Fig. 1D-F, Fig S1C&D)**. This increased production of TNFα **(Fig. 1D-F)** was significantly reduced by VNS. Additionally, R848-induced lung neutrophil recruitment was abrogated in mice subject to VNS (**Fig. 1G&H).**

**Figure 1.**
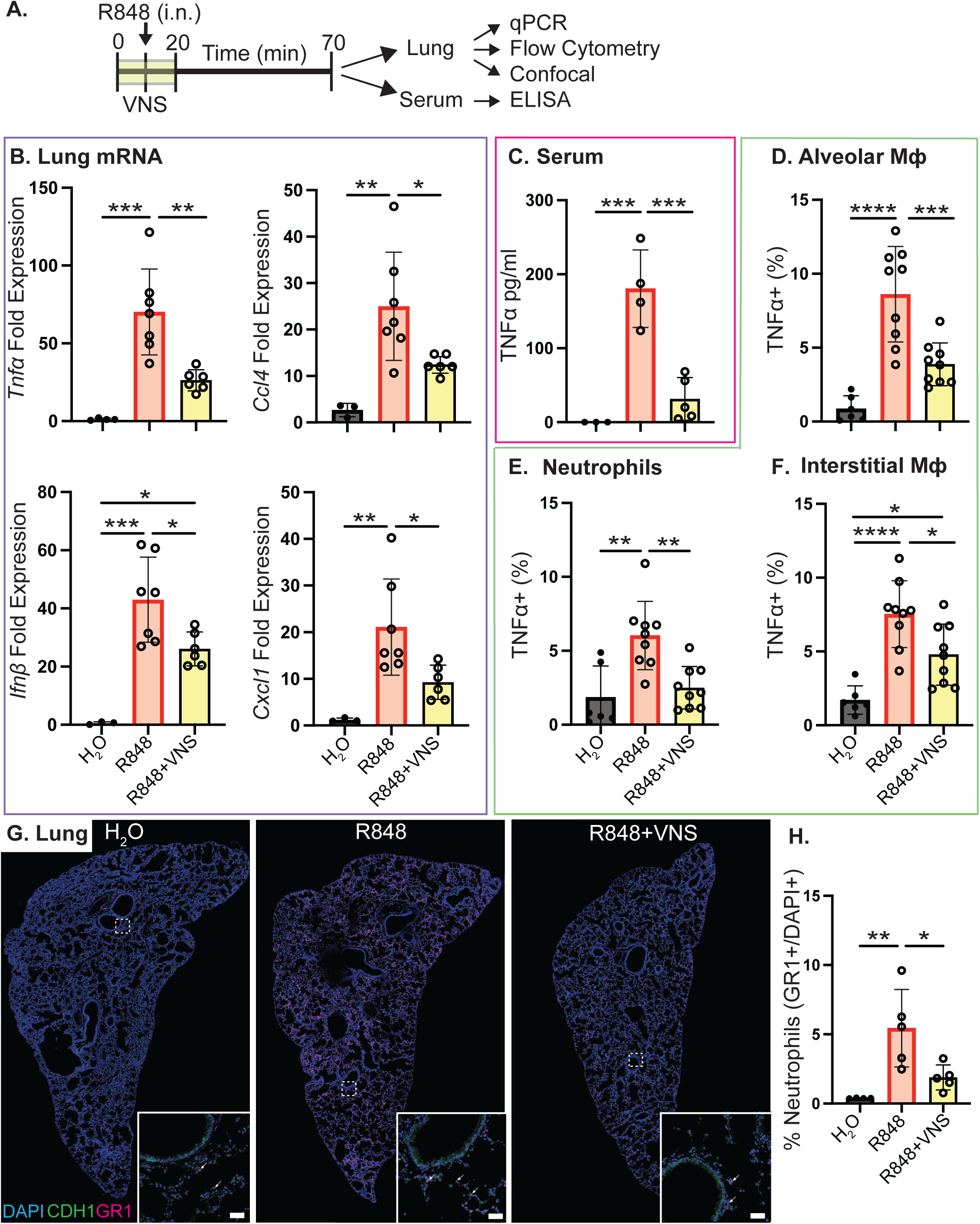
Electrical stimulation of the vagus nerve reduces TLR7-induced neutrophil and macrophage activation. **(A)** Experimental timeline. The vagus nerve was surgically isolated and electrically stimulated for 20 minutes. 0.25 mg/kg of R848 was instilled intranasally (i.n) 10 minutes post-VNS, and one-hour post-R848 challenge mice were euthanized. **(B)** mRNA expression of *Tnfα*, *Ccl4*, *Ifnp*, and *Cxcl1* in the right lung cranial lobe by qRT-PCR. **(C)** TNFα protein levels in the serum by ELISA. Flow cytometric analysis of TNFα+ **(D)** alveolar macrophages, **(E)** neutrophils, and **(F)** interstitial macrophages post-total lung digestion. **(G)** Immunofluorescence staining of neutrophils (GR1), epithelial cells (CDH1), and cell nuclei (DAPI) in the left lung lobe. Scale bar is 50µm. **(H)** Quantification of **(G)** as the percentage of GR1+ neutrophils in each lung section. Data represented as mean ± standard deviation (SD). One-way ANOVA was used for statistical analysis followed by post-hoc analysis with Tukey’s multiple comparisons test. *p< 0.05; **p<0.01; ***p<0.001; ****p<0.0001.

We devised a multiple-dose stimulation regimen to determine if VNS remained effective in subjects with prior inflammation. Mice were treated with three instillations of high dose R848 (2.5 mg/kg), followed by VNS and R848 restimulation **(Fig 2A)**. Prior R848 exposure with restimulation significantly increased TNFα production by neutrophils and alveolar macrophages in the lung compared to mice pretreated with water and stimulated with a single dose of R848 **(Fig S1 E&F)**. Protein concentration in the BALF was significantly increased with a single acute exposure to R848, and was further increased in mice pre-treated with R848 **(Fig. S1G)**. Having established R848 pre-stimulation increased lung inflammation compared to acute exposure, separate cohorts of mice were subjected to R848 pre-treatment followed by sham or VNS. As indicated, *Tnf*α mRNA in the lung **(Fig. 2B)**, and serum TNFα **(Fig. 2C)** were significantly reduced in mice that received VNS compared to sham stimulation.

**Figure 2.**
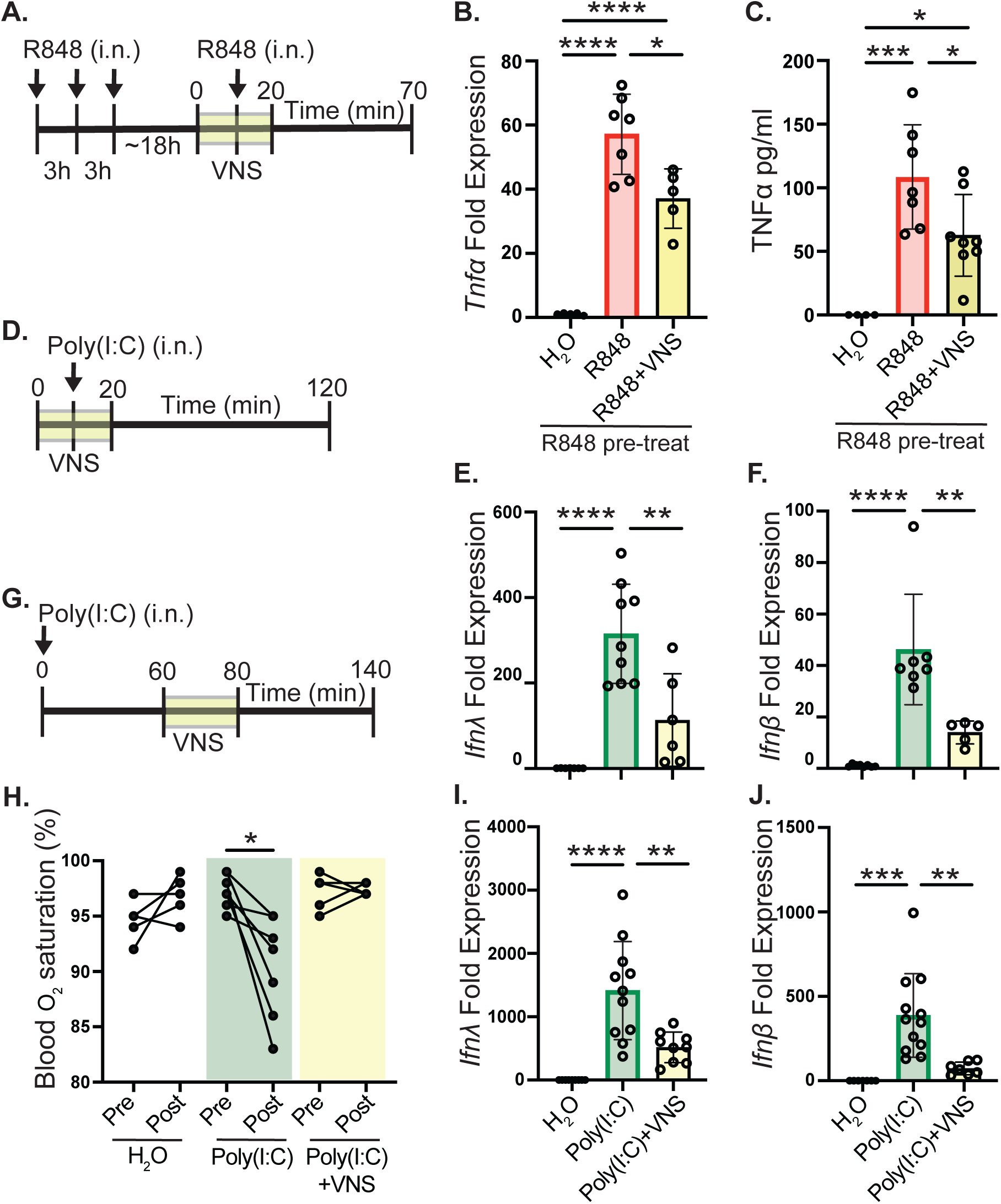
Electrical stimulation of the vagus nerve reduced both TLR7- and TLR3-induced lung acute and chronic inflammation. **(A)** Experimental timeline. High doses of 2.5 mg/kg R848 was instilled intranasally 3 times with 3-hour rest periods in between. After resting overnight, mice underwent 20-minute electrical VNS and intranasal instillation with 2.5 mg/kg R848. **(B)** mRNA expression of *Tnfα* in the right lung cranial lobe by qRT-PCR. **(C)** TNFα protein levels in the serum by ELISA. **(D)** Experimental timeline. Electrical VNS was performed for 20 minutes. 2.5 mg/kg of Poly(I:C) was instilled intranasally 10 minutes post-VNS, and two hours post-Poly(I:C) challenge mice were euthanized. mRNA expression of **(E)** *Ιfnλ* and **(F)** *Ιfnp* in the right lung cranial lobe by qRT-PCR. **(G)** Experimental timeline. 2.5 mg/kg Poly(I:C) was instilled intranasally and mice underwent 20-minute electrical VNS one hour post-Poly(I:C) challenge. Mice were euthanized one hour post-VNS. **(H)** Blood oxygen saturation was measured at 0- and 140-minute timepoints. mRNA expression of **(I)** *Ιfnλ* and **(J)** *Ιfnp* in the right lung cranial lobe by qRT-PCR. Data represented as mean ± standard deviation (SD). One-way ANOVA was used for statistical analysis followed by post-hoc analysis with Tukey’s multiple comparisons test. *p< 0.05; **p<0.01; ***p<0.001; ****p<0.0001.

We also assessed if VNS could reduce inflammation induced by the TLR3 agonist Poly(I:C) in both an acute **(Fig. 2D)** and multi-hour treatment regimen **(Fig. 2G)**. Indicating that VNS is broadly protective during acute lung inflammation, VNS, but not sham stimulation, significantly reduced *Ifnβ* **(Fig. 2E)**, and *Ifnλ* **(Fig. 2F)** induced by Poly(I:C) (i.n. 2.5 mg/kg) treated mice. To investigate the effect of VNS in treating induced lung inflammation, Poly(I:C) was administered 1h before VNS or sham stimulation and assessed after 2h **(Fig 2G)**. Poly(I:C) significantly reduced blood oxygenation **(Fig 2H),** and as expected increased *Ifnβ*, and *Ifn*λ expression **(Fig 2I&J).** Mice treated with VNS, but not sham stimulation 1h after Poly(I:C), had blood oxygenation concentrations equivalent to vehicle control mice, as well as significantly reduced *Ifnβ*, and *Ifn*λ expression **(Fig 2I&J)**. These data demonstrate that VNS can prevent and reduce inflammation and the resulting physiological changes.

To assess if these anti-inflammatory effects in the lung could be attributed to activation of the CAIP, we assessed the role of ChAT+ T-cells in our model. Using a ChAT-GFP reporter mouse, ChAT+ T-cells in the lung were found to be rare by flow cytometry **(Fig S2A).** We next performed VNS in WT (LCK.Cre^-^ ChAT^f/f^) and ChAT T-cell conditional knockout (LCK.Cre^+^ ChAT^f/f^) mice, of which the conditional knockout was confirmed (**Fig S2B&C)**. VNS significantly reduced R848-induced inflammation in WT and ChAT T-cell cKO mice **(Fig. S2D)**, demonstrating that VNS can reduce acute lung inflammation through a mechanism distinct from the CAIP. Although the hypothalamus-pituitary-adrenal axis activation and the release of corticosterone is well appreciated to be anti-inflammatory^26^, VNS did not increase serum corticosterone **(Fig. S2E)**. Moreover, we found that VNS did not induce the expression of *Il-10* mRNA in the lung **(Fig S2F)**. These data demonstrate that VNS activated an anti-inflammatory reflex that reduced acute lung inflammation by decreasing innate immune cell activation and recruitment.

### Stimulation of vagal afferent neurons reduces TLR7-induced lung inflammation

As discrete neuroimmune circuits are activated by selective vagal afferent or efferent stimulation^27^, we sought to determine the role of afferent-selective vagal nerve stimulation in the control of acute lung inflammation. We performed optogenetic activation of vagal ganglion before R848 treatment **(Fig 3A),** in VGlut2.Cre+ LSL-ChR2^YFP^ mice that conditionally express the activating channelrhodopsin (ChR2^YFP^) in vagal ganglia **(Fig. 3B)**. Selective activation of vagal ganglion by optical stimulation (465 nm) of VGlut2.Cre+ CHR2^YFP^ mice but not VGlut2.Cre+ eHR3^YFP^ (control) significantly reduced R848-induced serum TNFα **(Fig. 3C & Fig S3A)**. Additionally, R848-induced for *Tnf*α*, Ifn*β*, Cxcl1,* and *Ccl4* mRNA expression in the lung was significantly reduced upon optogenetic stimulation only in VGlut2.Cre+ ChR2^YFP^ **(Fig. 3D),** but not control VGlut2.Cre+ eHR3^YFP^ mice **(Fig S3B).** Optogenetic stimulation of the right vagal ganglion increased neuronal activity in the NTS and Area Postrema (AP) within the brainstem **(Fig 3E)**, loci that are known to receive input from the nodose ganglion^28^. Similarly, optogenetic activation of vagal ganglion of Phox2b.Cre+ LSL-ChR2^YFP^ mice significantly reduced R848-induced *Tnfα* and *Ifnβ* mRNA expression in the lung **(Fig. S3C)** and increased neuronal activity in the NTS and RVLM **(Fig. S3D)**, indicating that nodose ganglia activation was sufficient to reduce TLR7-induced lung inflammation.

**Figure 3.**
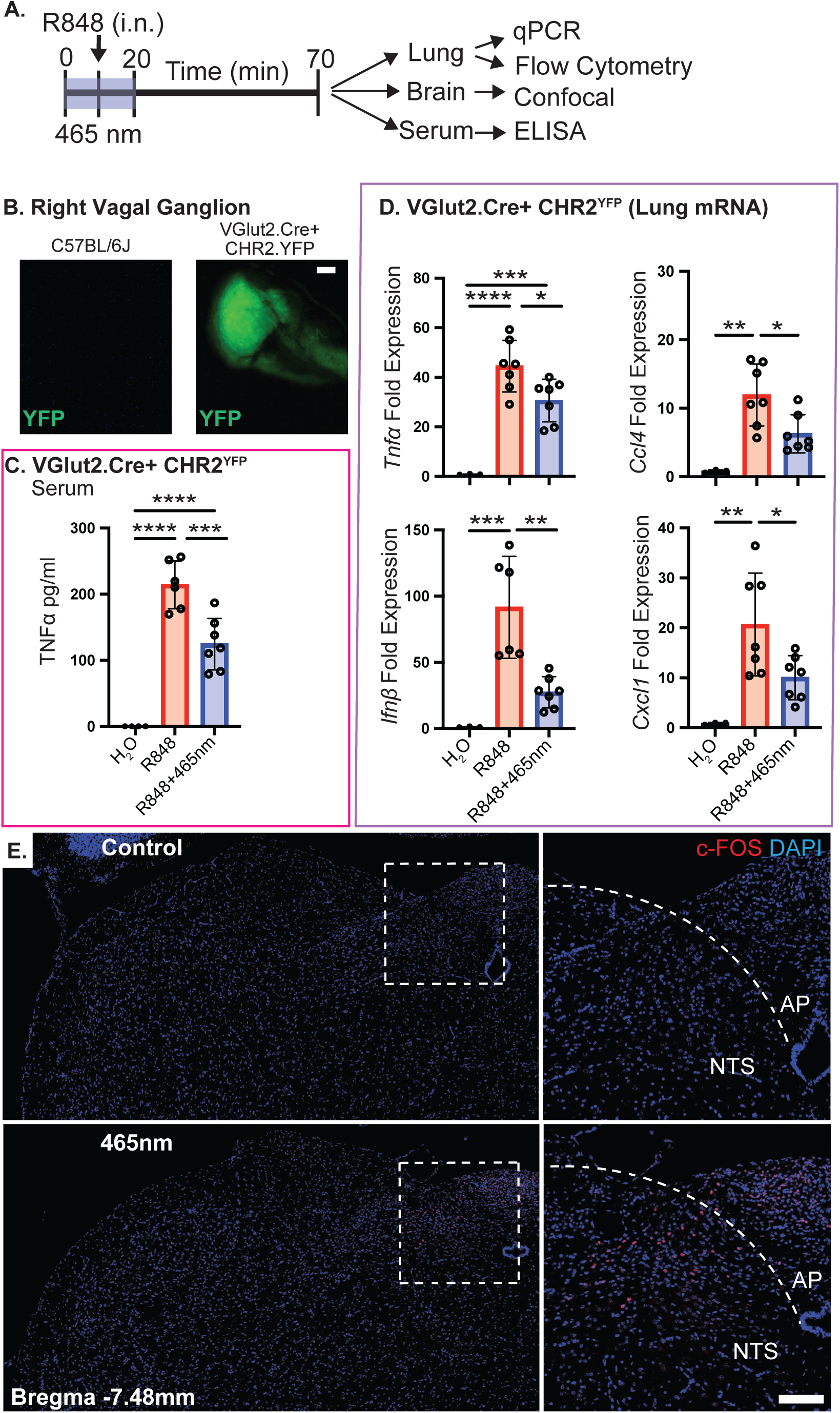
Optogenetic stimulation of vagal afferent neurons ameliorates TLR7-induced lung inflammation. **(A)** Experimental timeline. The nodose ganglion is carefully exposed (confirmed by existence of fluorescent reporter) and stimulated at 465 nm for 20 minutes. R848 was instilled 10 minutes into 465 nm stimulation, and mice were euthanized 1-hour post-R848 challenge. **(B)** Fluorescent YFP reporter localized in right nodose ganglion in wild-type (C57BL/6J) vs. transgenic optogenetic (VGlut2.Cre+ CHR2^YFP^) mice. Scale bar is 150 µm. **(C)** Serum TNFa protein detected by ELISA in VGlut2.Cre+ CHR2^YFP^ mice. mRNA expression of **(D)** *Tnf*α*, Ifn*β*, Ccl4, and Cxcl1* from right lung cranial lobe in VGlut2.Cre CHR2^YFP^ (experimental) mice by qRT-PCR. **(E)** Activated (c-FOS+DAPI+) neurons in the nucleus tractus solitarius (NTS), the brain region where vagal afferent neurons terminate, in stimulated control C57BL/6J mice vs. VGlut2.Cre CHR2^YFP^ mice. Scale bar is 100 µm. Data represented as mean ± standard deviation (SD). One-way ANOVA was used for statistical analysis followed by post-hoc analysis with Tukey’s multiple comparisons test. *p< 0.05; **p<0.01; ***p<0.001; ****p<0.0001.

### Control of acute lung inflammation by vagus nerve stimulation requires adrenal gland-derived epinephrine release

As activation of neuroimmune circuits culminates in catecholamine release in secondary lymphoid organs^29^, we assessed if VNS could lead to increased catecholamines. Electrical VNS, or selective optogenetic activation of the right vagal ganglion induced significantly increased epinephrine in the BALF and serum **(Fig. 4A&B)**. To evaluate whether VNS-induced epinephrine release contributed to control of acute lung inflammation, we first assessed if pharmacological inhibition of phenylethanolamine N-methyltransferase (PNMT) reduced epinephrine production. Treatment with the PNMT inhibitor (LY78335), but not the vehicle control, prevented VNS-induced increases in BALF and serum epinephrine, validating the efficacy of this inhibitor **(Fig S4A)**. In mice treated with the PNMT inhibitor, but not vehicle, VNS was no longer effective in attenuating R848-induced *Tnf*α and *Ifn*β mRNA expression **(Fig. 4C)**. As the adrenal glands are the primary source of peripheral epinephrine, we assessed the contribution of these glands in VNS-induced control of acute lung inflammation. While VNS reduced acute lung inflammation in sham-operated mice, protection afforded by VNS was lost in mice subject to adrenalectomy (ADX) **(Fig. 4D)**. Highlighting the importance of the adrenal glands, the ability of optogenetic stimulation of the right vagal ganglia to reduce R848-induced acute lung inflammation was significantly reduced in adrenalectomized mice **(Fig. 4E)**.

**Figure 4.**
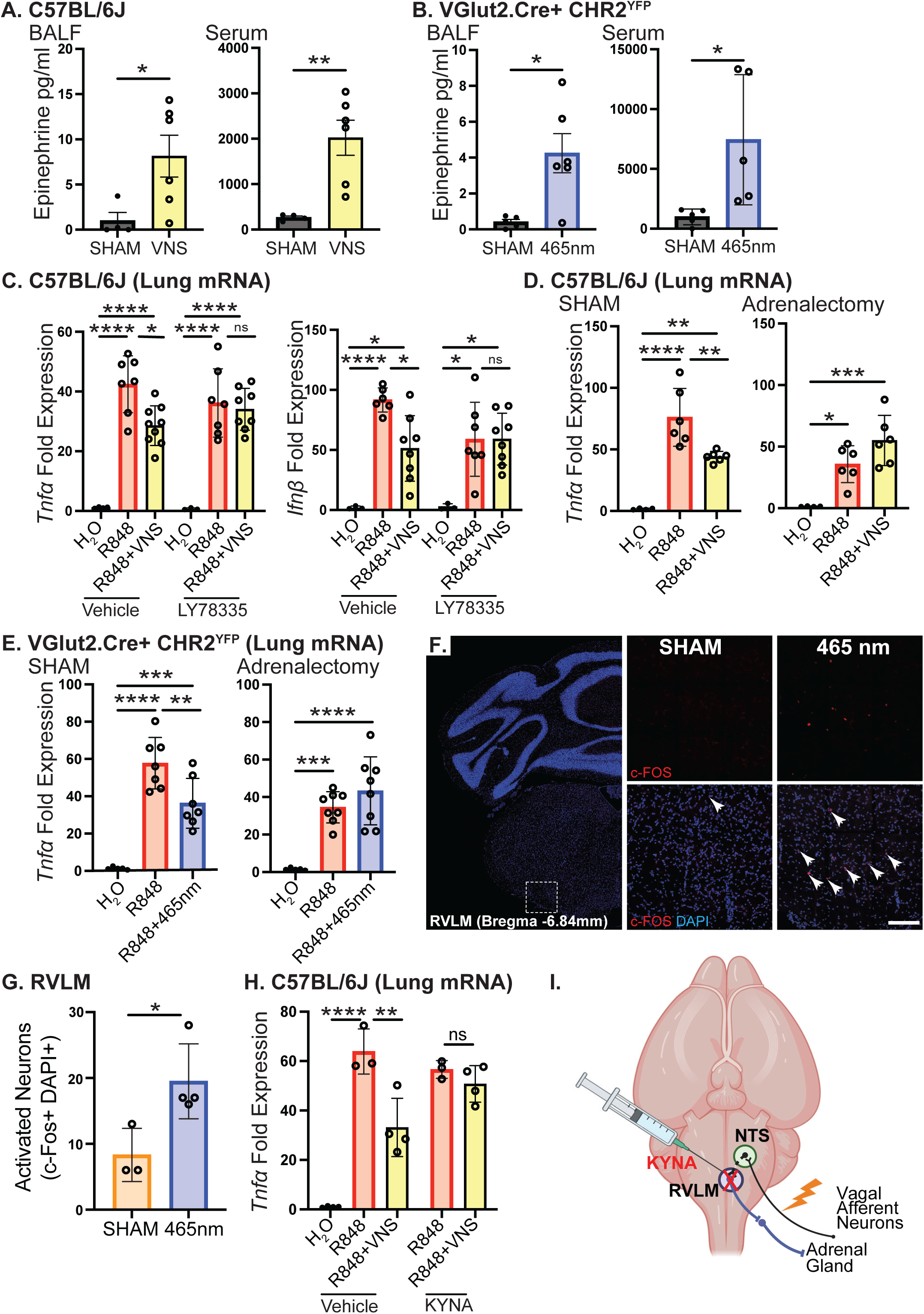
Vagus nerve activation induces adrenal gland-derived epinephrine release. Epinephrine in BALF and serum 5 minutes post **(A)** electrical VNS and **(B)** optogenetic activation of vagal afferent neurons detected by ELISA. **(C)** Right lung cranial lobe mRNA expression of *Tnf*a and *Ifn*b by qRT-PCR in mice challenged with i.p. delivery of water (vehicle) or PNMT inhibitor (LY78335) 1 hour prior to VNS. **(D)** Right lung cranial lobe mRNA expression of *Tnf*a in mice exposed to SHAM surgery (right) or adrenalectomy (left) 20 minutes prior to VNS. **(E)** Right lung cranial lobe mRNA expression of *Tnf*a in mice exposed to SHAM surgery (right) or adrenalectomy (left) 20 minutes prior to optogenetic stimulation of vagal afferent neurons. **(F)** Confocal image and **(G)** quantification of the RVLM in mice subject to 465 nm optogenetic or control (CTRL) stimulation at 590 nm. Neuronal activation was determined by colocalization of c-FOS and DAPI. A separate cohort of mice was used for this experiment. Scale bar is 100 µm. Data represented as mean ± standard deviation (SD). Student’s T-test or one-way ANOVA was used for statistical analysis followed by post-hoc analysis with Tukey’s multiple comparisons test. *p< 0.05; **p<0.01; ***p<0.001; ****p<0.0001.

VNS-stimulated release of epinephrine from the adrenal glands suggests the activation of the sympathetic nervous system. To determine the consequences of VNS in the brain, we assessed neuronal activation indicated by expression of the early immediate gene cFOS in the brain post electrical VNS, or selective optogenetic stimulation of the vagal ganglion. As expected, stimulation with either intact VNS, or optogenetic activation, increased neuronal activation (DAPI+ cFOS+) in the NTS **(Fig. S4B-D)**. Increased neuronal activation also occurred in mice that received either VNS **(Fig S4E&F)** or vagal ganglia selective optogenetic stimulation in the rostral ventrolateral medulla (RVLM), a sympathetic brain region **(Fig. 4F&G)**. To assess if neuronal activation of the RVLM was required as part of the neuroimmune circuit activated by VNS, we performed pharmacological blockade of glutamate receptors by stereotactic injection into this brain region. Mice receiving injection of Kynurenic acid into the RVLM and VNS were no longer protected from R848 induced *Tnfα* expression unlike vehicle and VNS-treated mice **(Fig 4H)**. Together, these data suggested that stimulation of vagal afferent neurons controls acute lung inflammation by activating the RVLM to induce epinephrine release from the adrenal glands **(Fig 4I)**.

### VNS attenuates R848-induced lung inflammation via β_2_AR activation

Epinephrine is a potent agonist of the β_2_AR, and the anti-inflammatory effects of VNS are dependent on β_2_AR activation in the spleen^16^. To determine if VNS-induced control of acute lung inflammation was dependent on the β_2_AR, we assessed the response of WT and β_2_AR KO mice. While VNS significantly reduced R848-induced serum TNFα **(Fig 5A)** and mRNA expression of *Tnf*α **(Fig 5B)** and *Ifn*β **(Fig S5A)** in the lungs of WT mice, this protective effect was lost in β_2_AR KO mice **(Fig. 5A-B, S5A)**. Further corroborating that the anti-inflammatory effect of VNS is dependent on the β_2_AR, VNS significantly reduced serum TNFα and mRNA cytokine transcripts in the lung of WT mice treated with vehicle but not the selective β_2_AR antagonist ICI 118 551 **(Fig. 5C&D, S5B)**. These data demonstrate that control of acute lung inflammation by VNS requires the β_2_AR *in vivo*.

**Figure 5.**
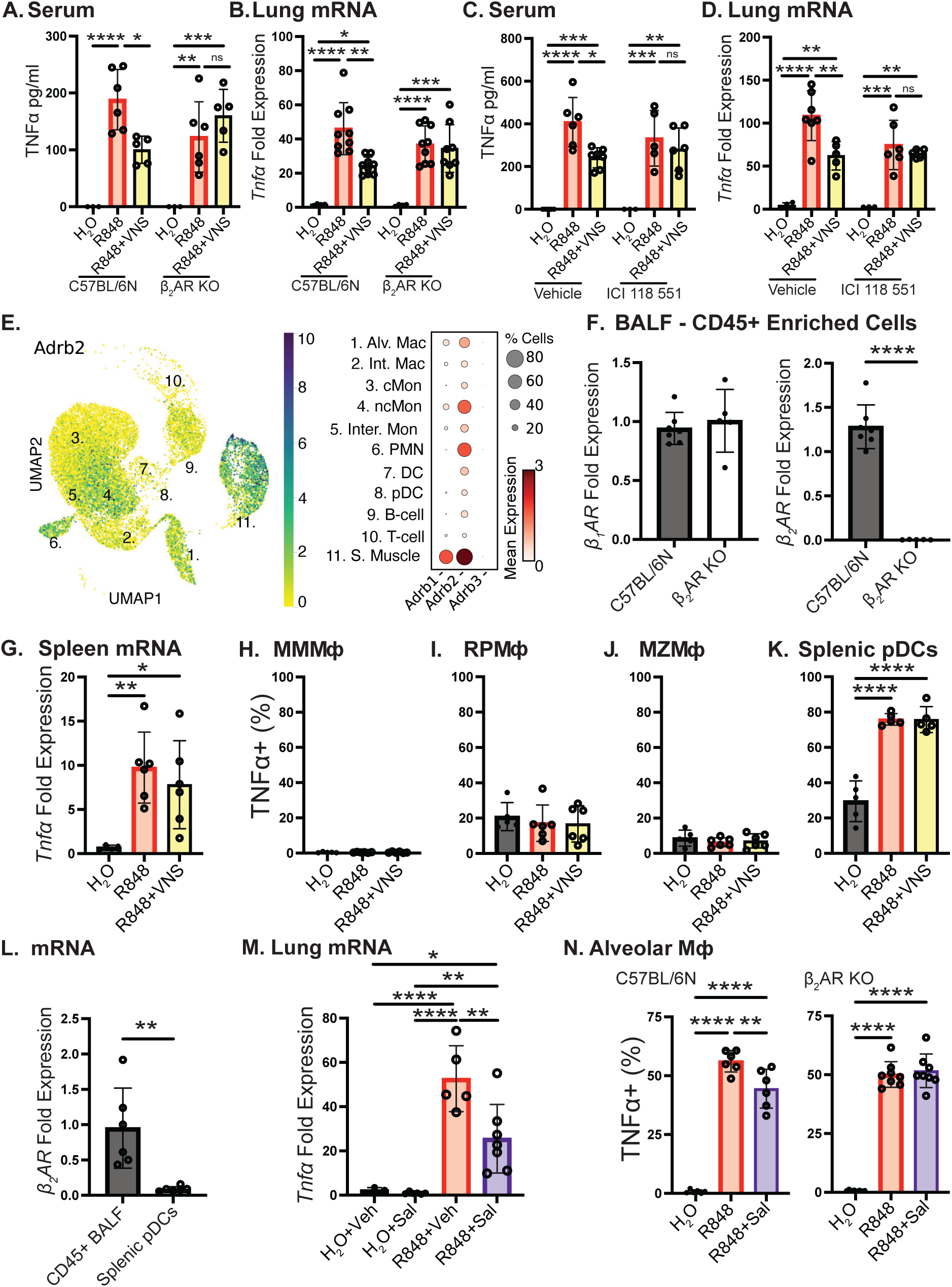
VNS requires β_2_AR activation for anti-inflammatory effects in lung. **(A)** Serum TNFα protein detected by ELISA and **(B)** lung mRNA expression of *Tnf*α by qRT-PCR in wild-type (C57BL/6N) and β_2_AR knockout mice subjected to electrical VNS. **(C)** Serum TNFα protein detected by ELISA and **(D)** lung mRNA expression of *Tnf*α in mice subjected to i.v. administration of vehicle (water) or β_2_AR antagonist (ICI 118 551), prior to challenge with R848 and electrical VNS. **(E)** Uniform manifold approximation and projection (UMAP) plot of all lung cells that express the *adrb2*, with the highest expression indicated by yellow. Expression of *adrb1*, *adbr2* and *adrb3* and percent of cells that expressed these genes of interest were evaluated. Populations with higher expression of the β_2_AR are indicated by larger circles and deeper coloring. **(F)** mRNA expression of β_1_AR (left) and β_2_AR (right) by qRT-PCR from CD45+ enriched BALF cells in wild-type (C57BL/6N) and β_2_ARKO mice. **(G)** mRNA expression of splenic *Tnf*α by qRT-PCR. Frequency of TNF+ **(H)** splenic marginal metallophilic macrophages (MMMΦ) **(I)** red pulp macrophages (RPMΦ) **(J)** marginal zone macrophages (MZMΦ) and **(K)** plasmacytoid dendritic cells (pDCs) by flow cytometry analysis. **(L)** mRNA expression of the β_2_AR on CD45+ enriched BALF cells and FACS-sorted splenic pDC’s by qRT-PCR. **(M)** mRNA expression of right lung cranial lobe *Tnf*α in mice treated with vehicle (water) or the β_2_AR agonist Salbutamol (Sal, 1mg/kg, i.v.) prior to challenge with water or R848. **(N)** Frequency of TNF+ alveolar macrophages treated *in vitro* with R848 +/-Salbutamol, isolated from the BALF of wild-type (C57BL/6N) or β_2_AR KO mice. Data represented as mean ± standard deviation (SD). One-way ANOVA was used for statistical analysis followed by post-hoc analysis with Tukey’s multiple comparisons test. *p< 0.05; **p<0.01; ***p<0.001; ****p<0.0001.

Expression of the β_2_AR has been reported on a multitude of immune cell populations. Analysis of public single-cell RNA sequencing data from the lung highlights this widespread expression, with β_2_AR transcripts identified on several immune cell populations including alveolar and interstitial macrophages, neutrophils, and monocytes. As expected, high levels of β_2_AR expression were also observed in smooth muscle **(Fig. 5E)**. Expression of β_1_AR was similar, and few of these cell types expressed β_3_AR RNA. Analysis of positively enriched CD45+ BALF cells, which are known to contain a high percentage of alveolar macrophages^30^, revealed β_1_AR and β_2_AR in WT mice and confirmed no increased β_1_AR and the absence of expression in β_2_AR KO mice **(Fig. 5F)**. Intranasal instillation of R848 also induced *Tnf*α expression in the spleen, and this was not reduced by VNS **(Fig. 5G)**. Intracellular cytokine staining and flow cytometry conducted on splenocytes revealed that R848 induced TNFα production was only observed in splenic plasmacytoid dendritic cells (pDC) (CD45+ B220+ CD317^high^) but not Marginal zone (CD19-, CD163-, F480+/-, Siglec1-, CD209b+), metallophilic (CD19-, CD163-, F480+/-, CD209b-, Siglec1+,), or red pulp (CD19-, F480+, CD163+) macrophages **(Fig. 5H-K**; **S5C**&**D**; **gating strategies)**. We reasoned that pDCs may lack the β_2_AR and assessed this by qRT-PCR performed on FACS-sorted pDCs. Compared to CD45+ enriched cells in the BALF, pDCs had negligible β_2_AR expression **(Fig. 5L)**.

Further demonstrating the anti-inflammatory role of the β_2_AR in lung inflammation, mice treated with the selective and efficacious β_2_AR agonist salbutamol (1 mg/kg, i.v.), but not vehicle-treated mice, had significantly reduced *Tnfα* expression in the lung, but not the spleen **(Fig. 5M, S5E)**. To assess the effects of β_2_AR activation on alveolar macrophages, BALF was harvested from WT and β_2_AR KO mice and stimulated with vehicle or salbutamol (1 μM), and with or without R848 (1 μg/mL). Intracellular cytokine staining demonstrated that R848 increased TNFα production in alveolar macrophages that was subsequently significantly inhibited by salbutamol only in WT but not β_2_AR KO mice **(Fig. 5N)**. These data demonstrate that innate immune cells in the lung, within the interstitial and alveolar space express β_2_AR, and that exogenous activation of this receptor is sufficient to block acute TLR7-induced lung inflammation.

## Discussion

Inappropriate immune cell activation in the lung and the ensuing inflammation that can occur in response to a variety of stimuli such as pathogens or ventilator induced injury are a significant health detriment. In this context, the ability to limit inflammation without reducing host protective responses is of significant interest. Communication between the nervous and immune systems has emerged as a mechanism to restrain immune cell activation and inflammation in a variety of organs systems including the lung. As the lung is highly innervated with sensory, parasympathetic and sympathetic neurons, it is not surprising that each of these has been implicated not only in the control of discrete aspects of lung physiology, but in host defense and the regulation of inflammation as well.

As part of this innervation, the vagus nerve provides both sensory afferent and efferent signaling within the lung^31^. Although the vagus nerve has a well-established immunoregulatory capacity in the spleen and lymph nodes during systemic inflammation, its role in lung inflammation is unclear. Our studies demonstrate that intact or selective vagal ganglion stimulation reduces TLR3 and TLR7 agonist-induced lung inflammation. We have also shown that VNS can attenuate inflammation in mice with prior, or ongoing inflammation. These data are in keeping with prior literature demonstrating that VNS can reduce inflammation and development of acute respiratory distress syndrome in mice challenged with LPS^32^, ventilator-induced lung injury^33^, or during influenza infection^21^. Critically, the ability of VNS to reduce lung inflammation occurred in a manner that is independent of the CAIP, as reduced proinflammatory cytokine expression was observed in ChAT T-cell conditional KO mice. As CD4+ T-cells that express this enzyme and release ACh are a required component of the CAIP, protection in the absence of these cells demonstrates a CAIP-independent pathway.

Corroborating these data and our interpretation, systemic inflammation induced by intravenous (i.v.) LPS was demonstrated to be controlled by the CAIP, and this pathway was without effect in the lung^34^. This lack of protection with VNS may be a consequence of the lung functioning as a unique neuroimmune circuit. We and others have previously shown that control of LPS-induced inflammation in the spleen can occur through two discrete pathways: one activated by vagal afferents, and the other activated by vagal efferents^16,29^. Our identification of a lung anti-inflammatory pathway that is activated by stimulation of vagal afferent as opposed to efferent neurons suggests the existence of several overlapping neuroimmune control pathways, capable of regulating immune cell activation across multiple tissues. A multitude of cell types in various organ systems including the lung have been reported to be regulated by neuroimmune communication, especially in the context of allergic asthma and acute respiratory distress syndrome^35–37^. Clinically, these conditions continue to have a high rate of mortality despite advances in acute care and ventilator technologies^38^. The ability to target innate immune cells and reduce inappropriate or overly exuberant activation, without generalized immune suppression is highly desirable.

Immune cells in the lung are naturally endowed with the ability to sense not only the presence of pathogens and host tissue damage, but also to receive neuronal signals by expression of neurotransmitter receptors^39^. Our data demonstrates that VNS and selective activation of vagal afferent neurons increased epinephrine release from the adrenal glands into the serum and within the alveolar space of the lungs. We further demonstrated that the adrenal glands, as the predominant source of epinephrine in the periphery^40^, are a critical part of a VNS-induced lung anti-inflammatory pathway as no protective effect of VNS was observed in adrenalectomized mice. Adrenal medullary chromaffin cells produce epinephrine by PNMT-mediated conversion of norepinephrine, a process that can be blocked by pharmacological inhibition^41,42^. Confirming that VNS regulation of acute lung inflammation required epinephrine, mice treated with a regimen of a PNMT inhibitor that prevented VNS-induced increases in epinephrine, did not have reduced lung inflammation despite VNS treatment. Although adrenal cortex derived corticosterone is a well-established negative regulator of inflammation^43^, we noted no VNS-induced increase of this steroid hormone in the serum. Together, these results suggest a unique pathway whereby activation of vagal afferents in the right cervical vagus nerve results in activation of sympathetic activity and release of catecholamines, but not activation of the hypothalamic-pituitary adrenal axis to regulate lung inflammation.

We identified that right cervical VNS and selective vagal ganglion stimulation induced neuronal activation (cFOS+ DAPI+) in discrete brain regions. VNS induced neuronal activation in the NTS, where afferent vagal fibers project, the Paraventricular nucleus of the hypothalamus (PVH), and the RVLM, a major sympathetic control center. While activation of the NTS can be anti-inflammatory in itself ^12,44^, the RVLM is a critical node in this new lung-protective neuroimmune circuit, as blocking glutamatergic signaling in this region prevented protection afforded by VNS. Although the NTS is generally considered to inhibit the RVLM, and consequently sympathetic activity^45^, glutamatergic projections from the NTS to the RVLM have been described^46^. These data suggest that selective activation of vagal afferents could induce peripheral sympathetic activity by activating this pathway, reducing inflammation.

This ability of vagal afferent activation to reduce peripheral inflammation during endotoxemia has also been observed to occur through increased splanchnic nerve activity^47^. Further corroborating this, VNS activation of neurons within the RVLM and subsequent sympathetic outflow can resolve inflammation in ischemia reperfusion injury (IRI) of the kidney^48^. Although reduced kidney IRI was independent of the adrenal glands, critically these studies targeted efferent vagal neurons. In this context, our data suggest that there are discrete neuroimmune pathways that can be activated by vagal efferent or afferent stimulation. It is an unresolved question how VNS is capable of eliciting a multitude of anti-inflammatory mechanisms in different organ systems. Undoubtedly, this reflects the complexity of the biology and context both of the neuronal stimulus delivered, and specific inflammatory insult being used. Vagal afferents are not a homogenous population as demonstrated not only by responsiveness to agonists or other stimuli, but also in terms of the projections within the NTS^49,50^. These projections of unique afferents to discrete NTS regions were proposed as an additional layer of specialization or spatially-based coding, as part of a mechanism where activation of select afferents would induce contextually relevant efferent signaling^51,52^. With this in mind, future studies defining precise NTS and RVLM neurons and circuitry could inform how VNS reduces inflammation in many models, and how to tune stimulation of these neurons to achieve better organ selectivity.

In assessing how the lung resident macrophages and recruited neutrophils could be regulated by epinephrine, we explored public scRNA sequencing data sets from mouse lung. While the β_2_AR in the CAIP has been established to respond to localized release of norepinephrine^53^, this receptor has a significantly higher affinity for epinephrine^54^. Our analysis of these datasets confirmed β_2_AR is expressed on many lung immune cell populations of interest including alveolar macrophages. Confirming that VNS mediated control of lung inflammation required β_2_AR, no protective effect of VNS was observed in mice deficient for this receptor, or mice receiving a highly efficacious and selective β_2_AR antagonist.

Demonstrating that the β_2_AR can reduce lung inflammation, administration of the selective agonist salbutamol significantly decreased pro-inflammatory cytokine expression. As evidence of an immune cell intrinsic effect of β_2_AR to attenuate R848-induced activation, alveolar macrophages stimulated with salbutamol prior to R848 produced significantly less TNFα production compared to R848 only controls. The expression and ability of β_2_AR to reduce TLR-induced inflammation is in keeping with prior literature^55–58^. Despite these prior reports of inhibition of TLR-induced inflammation or generation of an “alternatively activated” macrophage phenotype it is critical to note that, many of these studies use splenic, or macrophage-like cell lines^59^. Our assessment of lung resident macrophages further supports this anti-inflammatory role for the β_2_AR. The importance of cell specific responses is further highlighted in our data where although VNS abrogated R848-induced lung inflammation, no inhibition was observed in the spleen. Our analysis demonstrated that pDCs were the predominant source of TNFα in the spleen and that these cells do not respond to VNS most likely due to the absence of the β_2_AR. This then highlights a unique feature of the VNS-activated lung anti-inflammatory circuit, where activation of innate immune cells such as alveolar or interstitial macrophages, or neutrophils, is reduced, while other immune responses remain intact. These data suggest that selective activation of vagal afferent fibers could attenuate specific aspects of lung inflammation while allowing other host protective immune responses to develop.

## Methods

### Animals

Male and female C57BL/6J, C57BL/6N, B6J.129S6(FVB)-*Slc17a6^tm2(cre)Lowl^*/MwarJ (VGlut2.Cre); B6.129S-*Gt(ROSA)^26Sortm39(CAG-hop/EYFP)Hze^*/J (eHR3^YFP^) and B6.Cg-*Gt(ROSA)26Sor^tm32(CAG-COP4*H134R/EYFP)Hze^*/J (ChR2^YFP^) mice were originally purchased from Jackson Laboratories (Bar Harbor, ME, USA) and used to establish a breeding colony. Constitutive β_2_AR knock out (β_2_AR KO) mice were purchased from the Mouse Biology Program (University of California, Davis). Mice with a conditional KO for ChAT in T-cells (LCK.Cre^+^ ChAT^f/f^) were originally purchased from Jackson Laboratories and described previously ^60^. Male and female mice aged 8-12 weeks were utilized for all experiments. Every experimental intervention *in vivo* was compared to sham-treated subjects. All animals had *ad libitum* access to food and water, and all procedures were approved by the UC Davis Institutional Animal Care and Use Committee.

### Bronchoalveolar lavage

Mice were euthanized with CO_2_ and placed in a supine position. A small midline cervical incision was performed allowing separation of the sublingual glands, revealing the trachea. Lung lavage was performed using warmed BALF buffer (2 mM EDTA, 0.5% FBS, 1x PBS) with a 1mL syringe and 3/8-inch 26g needle attached. The needle was inserted between the cartilaginous rings of the trachea and a lavage was repeated 5x to retrieve the cells. Successful lavage was confirmed by lung inflation, and recovered fluid was kept on ice until further processing.

### BALF cell culture

BALF was extracted as described above. Cells were treated with Ack lysis solution for 1 minute at room temperature prior to excess washing. Cell were then resuspended in 1x DMEM +10% FBS + 1 mM sodium pyruvate + 10 mM HEPES + 1% Pen/Strep + 1X GolgiPlug. Cells were either incubated with H2O, 1 µg/mL R848, or 1 µg/mL with 1 µM Salbutamol for 1 h at 37°C. Cells then underwent flow cytometry staining and analysis for alveolar macrophages and TNFα expression.

### BALF Protein Concentration Determination

BALF was extracted as described above. Lavage fluid was centrifuged at 400 x g for 5 minutes and the supernatant was subjected to a Pierce BCA assay (Cat# A55864 Thermo Fisher Scientific) according to manufacturer’s instructions. Optical density was recorded at 562 nm using a plate reader, with sample concentrations determined by linear regression using a standard curve.

### Adrenalectomy (ADX)

Male and female mice were anesthetized using isoflurane, and body temperature was monitored and maintained at 37°C using a thermocouple-controlled heating pad (World Precision Instruments, Sarasota FL). Mice were placed in a supine position and a midline incision was made on the abdomen, allowing for gentle retraction of the intestinal tract, and removal of the right and left adrenal glands. The intestine and liver were placed back into position, and covered with sterile gauze moistened with warm, sterile saline. The mice remained under anesthesia for 20 minutes prior to beginning VNS.

### Vagal nerve stimulation

Male and female mice were anesthetized using 1-2% isoflurane and body temperature was maintained at 37°C using a thermocouple-controlled heating pad (World Precision Instruments). The right cervical vagus nerve was isolated, allowing careful placement on a bipolar hook electrode (FHC, Bowdoin, ME, USA). Mice were then subjected to 20 minutes of VNS (5 V, 2 ms, monophasic square wave, 5 Hz), or sham conditions, using a Grass stimulator S88 with a stimulus isolation module described in previous publications^61^. Resiquimod (R848, InvivoGen, tlrl-r848, 0.25 mg/kg in 10 μL as droplets) or Polyinosinic-polycytidylic acid (Poly(I:C), InvivoGen, tlrl-pic, 2.5 mg/kg in 60 μL as droplets) were administered by intranasal instillation after the first 10 minutes of stimulation. Isoflurane is maintained throughout with delivery by nosecone and animals resting prone at a slight incline. Serum was collected via a cardiac puncture one hour or two hours post-R848 or Poly(I:C) stimulation respectively. Mice were euthanized by cervical dislocation, and lung and spleen tissues were collected for flow cytometry or qRT-PCR. Alternatively, 2.5 mg/kg R848 was instilled intranasally three times with 3-hour rest periods in between. Mice were then returned to the vivarium overnight and underwent the treated as described above but with 2.5 mg/kg R848 where applicable. For Poly(I:C) pretreatment, mice were placed under 1-2% isoflurane for five minutes before blood oxygenation was measured MouseSTAT Jr (Kent Scientific, Torrington, CT) and 2.5 mg/kg Poly(I:C) was instilled intranasally. Mice were kept under isoflurane for one hour and underwent 20 minutes of vagal nerve stimulation as described above. Mice were kept under isoflurane for an additional hour and blood oxygenation was measured before euthanasia.

### Optogenetics

Male and female VGLUT2.Cre+ CHR2^YFP^ mice were anesthetized using 1-2% isoflurane. A small incision was made over the trachea and the sublingual glands were separated with a retractor. Additional retractors were placed carefully on surrounding muscles to reveal the right nodose ganglion, which was confirmed by visualization of the YFP reporter, and positioning of the fiberoptic cable. Stimulation with 465 nm, was achieved using a computer-controlled LED light source (Plexon Inc. Dallas, TX) for 20 minutes of stimulation (2 ms pulse duration, 10 Hz) was then performed. R848 was instilled after the first 10 minutes of stimulation. 1-hour post-R848 challenge, mice were euthanized, and tissues were collected as described above. To control for any heat emitted from the fiber optic probe, VGLUT2.Cre+ eHR3^YFP^ mice (activated at 590 nm) underwent the same experimental parameters outlined above, including stimulation at 465 nm.

### Pharmacological agents

Selective blockade of β_2_AR receptors was performed with ICI 118 551 (Tocris, Minneapolis, MN): A 0.5 mg/kg dose or vehicle control (water) was instilled as droplets intranasally 10 minutes prior to beginning VNS. To activate β_2_AR, Salbutamol hemisulfate was used (Tocris, Minneapolis, MN). A 1 mg/kg dose or vehicle (water) was i.v. injected 15 minutes prior to challenge with R848 (0.25 mg/kg). Inhibition of PNMT was performed with LY78335 (Santa Cruz Biotechnology). A 25 mg/kg dose or vehicle (water) was i.p. injected 60 minutes prior to beginning subsequent experiments.

### Quantitative real-time PCR

Gene expression in the lung was analyzed by real-time PCR (qRT-PCR). RNA was extracted by lysing lung tissue in TRIzol (Invitrogen) using 5 mm stainless steel beads in a bead beater (Qiagen), per manufacturer’s instructions. To synthesize cDNA, an iSCRIPT reverse transcriptase kit (Bio-Rad, Hercules, CA, USA) was used. qRT-PCR was then performed for the following targets using primer pairs sourced from Primerbank^62^. *Actb* forward 5’-GGCTGTATTCCCCTCCATCG-3′ reverse 5’-CCAGTTGGTAACAATGCCATGT-3’; *Tnf*a forward 5’-CCTCACACTCAGATCATCTTCT-3’ reverse 5’-GCTACGACGTGGGCTACAG-3’; *Cxcl1* forward 5’-TCCAGAGCTTGAAGGTGTTGCC-3’ reverse 5’-AACCAAGGGAGCTTCAGGGTCA-3’-*Ifn*b forward 5’-CAGCTCCAAGAAAGGACGAAC-3’ reverse 5’-GGCAGTGTAACTCTTCTGCAT-3’; *Ccl4* forward 5’-TTCCTGCTGTTTCTCTTACACCT-3’ reverse 5’-CTGTCTGCCTCTTTTGGTCAG-3’. *Ifnl* forward 5’-GTTCAAGTCTCTGTCCCCAAAA-3′ reverse 5’-GTGGGAACTGCACCTCATGT-3’. Amplification was conducted using a QuantStudio6 (Thermo Fisher Scientific, Waltham, MA, USA) machine. *Actb* was used as a reference gene for each sample with fold expression calculated by the 2^(-ΔΔCT)^ method.

### Serum cytokine analysis

To analyze the concentration of TNFα in the serum, a sandwich ELISA kit (Thermo Fisher Scientific) was used. Manufacturer’s instructions were followed but briefly, 96-well maxisorp microtiter plates (NUNC, Thermo Fisher Scientific) were coated with capture antibody and washed thoroughly with PBS-Tween20 0.5% v/v prior to adding serum samples for incubation. The plates were washed extensively again followed by a brief incubation with the biotin conjugated detection antibody. Streptavidin HRP was then added followed by a final wash to clear any unbound enzyme. Plates were developed by adding TMB substrate for 15 minutes before stopping the reaction with 1 N H_2_SO_4_. Optical density was recorded at 450 nm and 570 nm using a plate reader, with sample concentrations determined by linear regression using a standard curve.

### Catecholamine Analysis

To analyze the concentration of catecholamines in bronchoalveolar lavage fluid (BALF) and serum, a commercially available competitive ELISA kit was used (Rocky Mountain Diagnostic, Inc. CO, USA). Male and female mice were exposed to 5 minutes of control, electrical, or optogenetic stimulation. Catecholamines were stabilized in the serum and BALF with the addition of EDTA (final concentration 1 mM) and sodium metabisulfite (final concentration 4 mM) and assessed as per manufacturer instructions.

### Imaging

Brains were perfused and post-fixed in 4% PFA for 48 h, followed by incubation in 15% and 30% sucrose in PBS at 4°C for 24 h respectively. Brains were subsequently embedded in OCT medium (Fisher Scientific, Cat #23-730-571) and stored at −80°C overnight. Coronal sections (25 μm) were cut using a cryostat (Leica, Germany). Sections of NTS were collected between Bregma −7.48mm and −7.32 mm, while sections of RVLM were collected around Bregma −6.84 mm. After blocking in 5% BSA (w/v) and normal donkey serum, samples were incubated in primary antibody overnight at 4°C. Slides were washed extensively in TBS-tween20 0.1% v/v and incubated in secondary antibodies for 1 h at room temperature. Nuclei were counterstained with DAPI in 1xPBS, washed and mounted in Prolong gold (ThermoFisher, Waltham, MA). Neuronal activation was assessed by quantifying the numbers of cFos^+^/DAPI^+^ cells in the NTS and RVLM regions. Coronal sections were stained with rabbit anti-cFos (Cell Signaling, Cat# 2250S, 1:100), followed by donkey anti-rabbit Alexa Fluor 546 (Invitrogen, Cat# A10040, 1:200). Confocal images were taken with a 20x and 40x objective (SP8; Leica) with a 0.70-μm z-step size. The numbers of cFos^+^/DAPI^+^ cells were quantified within the defined regions using Imaris (Oxford instruments, UK).

Left lungs were collected and post-fixed in 4% PFA for 48 h, followed by incubation in 15% and 30% sucrose in PBS at 4°C for 24 h respectively. Lung samples were embedded in OCT medium (Fisher Scientific, Cat #23-730-571) and stored at −80°C overnight. Cryosections of lung tissues (10 μm) were used for immunofluorescence staining with the protocol described above. Cryosections were stained with the following primary antibodies: biotin mouse anti-Ly-6G (Gr-1) (Tonbo, Cat# 30-5931-U500, 1:100) and rabbit anti-E Cadherin (ECM Biosciences, Cat #CP1921, 1:400), followed by secondary antibodies: streptavidin Alexa Fluor 647 (Invitrogen, Cat #S32357, 1:200) and donkey anti-rabbit Alexa Fluor 546 (Invitrogen, Cat# A10040, 1:200). Confocal images were taken with a 20x and 40x objective (SP8; Leica) with a 0.70-μm z-step size. The numbers of GR1+ cells (neutrophils) were quantified with the following steps: We used cellpose 2 from the gui to segment the nuclei and generate a mask. The pretrained nuclei model with flow set to 1.5 and cell prob to −2 was used. The size of the nuclei was calibrated in cellpose 2 and the resulting mask as a tiff file. Cells that had a mean GR1 intensity higher than threshold were found using a customized python code and determined visually. Total GR1+ cells were divided by the total number of DAPI+ cells within the respective tissue sections.

### Stereotactic Injection

10- to 11-week old mice were placed under 1-2% of isoflurane for five minutes and immobilized in a stereotaxic apparatus. A single, 0.4-μl injection of either saline or kynurenic acid (KYNA, Tocris, 0223, 0.2 μM) was delivered over a 2-min period into the left RVLM (coordinates from bregma: −6.50 mm posterior, +1.50 mm lateral, and −6.5 mm ventral). Immediately after, another injection was delivered into the right RVLM (coordinates from bregma: −6.50 mm posterior, −1.50 mm lateral, and −6.5 mm ventral). The exposed skull was kept moist with a sterile gauze and saline. Mice was kept under isoflurane for another 10 minutes before VNS was performed as described above. One-hour post-R848 challenge, mice were euthanized by cervical dislocation and lung tissues were collected for qRT-PCR.

### Flow cytometry

Whole lungs were extracted from mice treated as described above and placed in a C-tube with 2.5 mL of 1x Buffer S, 100 μL of enzyme D, and 15 μL of enzyme A from the Mouse Lung Dissociation Kit (Cat# 130-095-927, Miltenyi Biotec, North Rhine-Westphalia, Germany). Whole spleen for macrophage and pDC analysis was digested using the same kit. Protocol was followed as described by manufacture until cells were strained through 100 μm cell strainer. Single cell suspension had red blood cells lysed using ACK lysis buffer for 5 minutes at room temperature before diluting with FACS buffer (PBS + 2% FBS). Cells were enumerated on a hemocytometer using trypan blue to stain dead cells. Single cell suspension is first incubated with Fc Shield (Cat# 553142 BD Biosciences, San Jose CA) for 25 minutes at 4°C before being incubated with antibodies to identify certain populations of interest for 30 minutes at 4°C. Viable cells were identified using Live/Dead Fixable Aqua Dead Cell Stain Kit (Cat# L34957 ThermoFisher, Waltham MA) or Ghost V510 (Cat# 13-0870-T100 Tonbo Biosciences, San Diego CA). For identifying alveolar and interstitial macrophages in the lung, cells were incubated with antibodies to CD45 efluor450 (Cat# 48-0451-82 Invitrogen, Waltham MA), CD11c PE-Cyanine7 (Cat# 60-0114-U100 Tonbo Biosciences), CD11b AF700 (Cat# 56-0112-82 Invitrogen, Waltham MA), CD64 PE (Cat# 130-118-684 Miltenyi Biotech), and SiglecF BV650 (Cat#740557 BD Biosciences, San Jose CA). For identifying neutrophils and monocytes in the lung, cells were incubated with antibodies to CD45 efluor450 (Cat# 48-0451-82 Invitrogen), MHCII PE-efluor610 (Cat# 61-5321-82 Invitrogen), CD11b AF700 (Cat# 56-0112-82 Invitrogen, Waltham MA), CD64 BV605 (Cat#139323 Biolegend), Ly6C PE-Cyanine7 (Cat# 25-5932-82 Invitrogen), and Ly6G FITC (Cat# 35-1276-U100 Tonbo Biosciences). For identifying T cells, B cells, and NK cells in the lung, cells were incubated with antibodies to CD45 AF488 (Cat#103122 Biolegend), CD4 AF700 (Cat#100430 Biolegend), CD8b APC (Cat# 17-0083-81 eBioscience), CD5 PE-Cy7 (Cat# 25-0051-81 Invitrogen), CD19 PE-efluor610 (Cat#61-0193-82 Invitrogen), and CD161 Biotin (Cat# 30-5941-U500 Tonbo Biosciences) identified with Streptavidin BV605 (Cat# 563260 BD Biosciences). For identifying splenic macrophages, single cell suspensions were incubated with antibodies to F4/80 PE (Cat# 12-4801-82 Invitrogen), CD163 APC (Cat# 17-1631-82 Invitrogen), CD209b AF488 (Cat# 53-2093-82 Invitrogen), CD169 AF700 (Cat# FAB5610N-100UG R&D Systems, Minneapolis, MN), and CD19 PE-efluor610 (Cat# 61-0193-82 Invitrogen). For identifying splenic pDC, cells were incubated with antibodies to CD45 efluor450 (Cat# 48-0451-82 Invitrogen), CD317 PE (Cat# 12-3172-82 Invitrogen) and B220 PerCP-Cyanine5.5 (Cat# 65-0452-U100 Tonbo Biosciences). Cells were then fixed and permeabilized using BD Cytofix/ Cytoperm (Cat# 51-2090KZ BD Biosciences) for 25 minutes at 4°C and washed using BD Perm/ Wash Buffer (Cat# 51-2091KZ BD Biosciences). Intracellular antibodies were diluted in BD Perm Buffer and incubated with the single cell suspension for 30 minutes at 4°C. Intracellular cytokines were identified using antibodies that bind to TNFα AF647 (Cat# 506314 Biolegend) or TNFα BV421 (Cat#563387 BD Biosciences). Cells were washed and resuspended in FACS buffer, with data acquisition performed on an LSR II (BD Biosciences). Data analysis was performed using FlowJo v10 (Ashland, OR: Becton Dickenson).

### Magnetic enrichment

To enrich CD45+ cells from BALF, a lung lavage was performed in euthanized male and female mice using warmed BALF buffer (2 mM EDTA, 0.5% FBS, 1x PBS). Collected cells were stored on ice prior to centrifugation (200g, 4°C). Cells were treated with ACK Lysis for 2 minutes, followed by dilution with FACS buffer. Cells were centrifuged again followed by a 15-minute incubation with Fc Shield (Cat# 553142 BD Biosciences, San Jose CA) on ice. A biotinylated CD45 antibody (Cat# 13-0451-82 Invitrogen) was added, and incubated for another 15 minutes on ice. The antibody was washed with FACS buffer, followed by application of Streptavidin particles plus (Cat# 557812, BD Biosciences). Cells were incubated with the Streptavidin particles for 30 minutes at 4°C on a rocker. Positive enrichment was then performed utilizing a cell separation magnet (Cat# 552311, BD Biosciences): 1 mL of FACS buffer was added to the cells and beads and placed on the magnet. Cells were allowed to bind for 8 minutes prior to removing the negative fraction. This was repeated 3x prior to preparing the cells for qRT-PCR.

### FACS sorting

Spleens were digested as described above. Prior to undergoing FACS sorting, pDC were positively enriched following the protocol described above for magnetic enrichment, using a biotinylated CD317 antibody (Cat# 13-3172-82, Invitrogen). Cells were then incubated and stained for 25 minutes with antibodies to identify viable pDC: Live/ Dead Fixable Aqua Dead Cell Stain Kit (Cat# L34957 ThermoFisher, Waltham MA), CD45 efluor450 (Cat# 48-0451-82 Invitrogen), B220 PerCP-Cyanine5.5 (Cat# 65-0452-U100 Tonbo Biosciences), and CD317 PE (Cat# 12-3172-82 Invitrogen). Cells were sorted into FACS buffer on an Astrios EQ prior to prepping for qRT-PCR.

### ChAT Validation

Spleens were excised from euthanized C57BL/6 or Cre-or Lck.Cre+ ChAT f/f mice and reduced to a single cell suspension. T and B cells were identified using anti-CD3 APC (Cat# 20-0031-U100 Tonbo Biosciences) and anti-B220 PerCP-Cy5.5 (Cat# 65-0452-U025 Tonbo Biosciences) and sorted on Sony MA900 Cell Sorter. Genomic DNA was then extracted by resuspending cells in digestion buffer (50mM KCL, 10mM Tris-HCl, 1% Triton-X, 2mM EDTA) and incubation at 56°C for 30 minutes then 95°C for 15 minutes. PCR primers were designed to detect ChAT genomic sequence with F: 5’-GGCCATTGTGAAGCGGTTTG-3’ which binds to exon 3 and R: 5’-CCGCCTCAGGACTCTTC-3’ which binds to the modified region with the loxP site after exon 4 and an expected band at 1.4kb. During loxP recombination in the presence of Cre, exon3 and 4 are excised, with no expected band. The same genomic DNA was also used for a PCR reaction to detect Cre using the Lck.Cre genotyping primers from Jackson labs, resulting in a Cre+ band at 300 bp and an internal control band at 200 bp.

### Re-analysis of publicly available datasets

Previously published and publicly available data from the Tabula Muris Senis project were downloaded from Figshare (https://figshare.com/projects/Tabula_Muris_Senis/64982) as .H5ad files, containing the original cell type annotation ^63^. This data set was visualized using Scanpy (v 1.9.1) in Jupyter notebook from a Python environment (v3.10.9), as either UMAP or dot plots for expression of specific genes of interest in the previously defined cell types, and the resulting plots were saved as SVG outputs.

## Acknowledgements

The authors would like to thank Dr. Nicole Baumgarth for her expertise and guidance. This work was a product of funding provided by the Chan-Zuckerberg Initiative CZI DAF2020-217656 (CR), NIH NIAID R56AI179656 (CR), and the “Animal Models of Infectious Disease Training Program” T32AI060555 (MC).

## Author Contributions

C.R., MG, and K.M. conceptualized experiments. KM executed electrical and optogenetic experiments and qRT-PCR. M.C. designed and executed cellular staining and flow cytometry. E.T. performed terminal stereotactic surgical procedures, electrical vagal nerve stimulation, immunostained and imaged lung and brain tissues. K.S. performed VNS and optogenetic experiments. S.S. optimized culturing and treatment of macrophages. I.B.M designed cell counting code. K.S., W.L., and J.A. performed *in vivo* salbutamol experiments. C.R. and K.M. wrote the paper. All authors were asked to edit the manuscript.

## Data Availability Statement

All data supporting the findings of this study are located within the paper and the supplementary information.

## Code Availability Statement

Previously published and publicly available data from the Tabula Muris Senis project were downloaded from Figshare (https://figshare.com/projects/Tabula_Muris_Senis/64982) as .H5ad files, containing the original cell type annotation ^63^.

## Supplementary Information

Supplementary Information is available for this paper.

---

**Figure S1.**
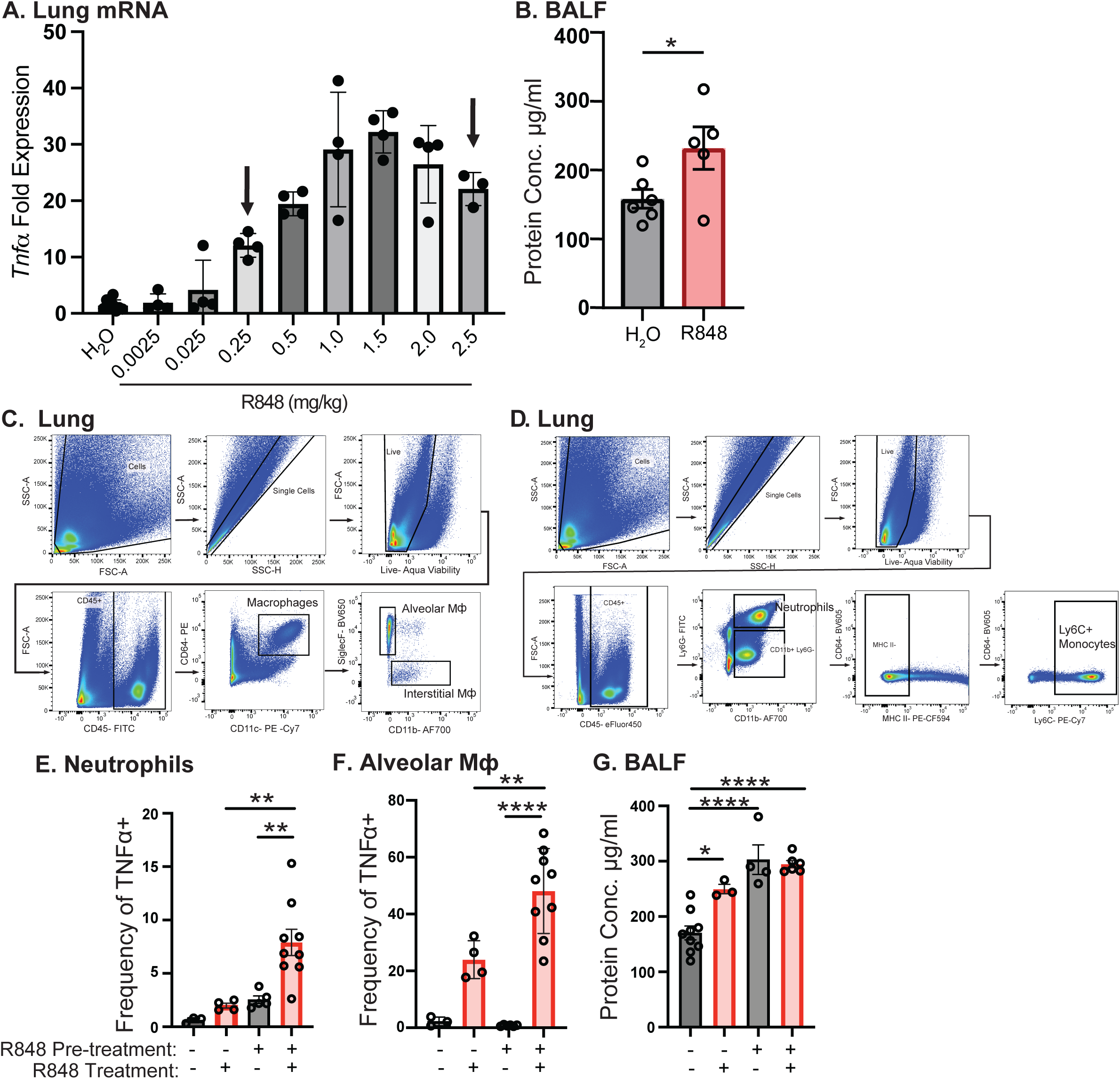
**(A)** 1-hour R848 (intranasal) dose response by mRNA expression of *Tnfα* in the right lung cranial lobe by qRT-PCR. Arrow points to doses used for experiments. **(B)** Protein concentration detected by ELISA in BALF 1-hour post R848 challenge. **(C)** Gating strategy used to distinguish alveolar and interstitial macrophages in the lung. **(D)** Gating strategy used to distinguish neutrophils and monocytes in the lung. Frequency of TNFα+ lung **(E)** neutrophils and **(F)** alveolar macrophages post challenge with 2.5 mg/kg R848, instilled intranasally 3 times with 3-hour rest periods in between. After resting overnight, mice were subject to a final intranasal instillation with 2.5 mg/kg R848. **(G)** Protein concentration detected in BALF following the dosing schema described in (E&F). Data represented as mean± standard deviation (SD). One-way ANOVA was used for statistical analysis followed by post-hoc analysis with Tukey’s multiple comparisons test. *p< 0.05; **p<0.01; ***p<0.001; ****p<0.0001.

**Figure S2.**
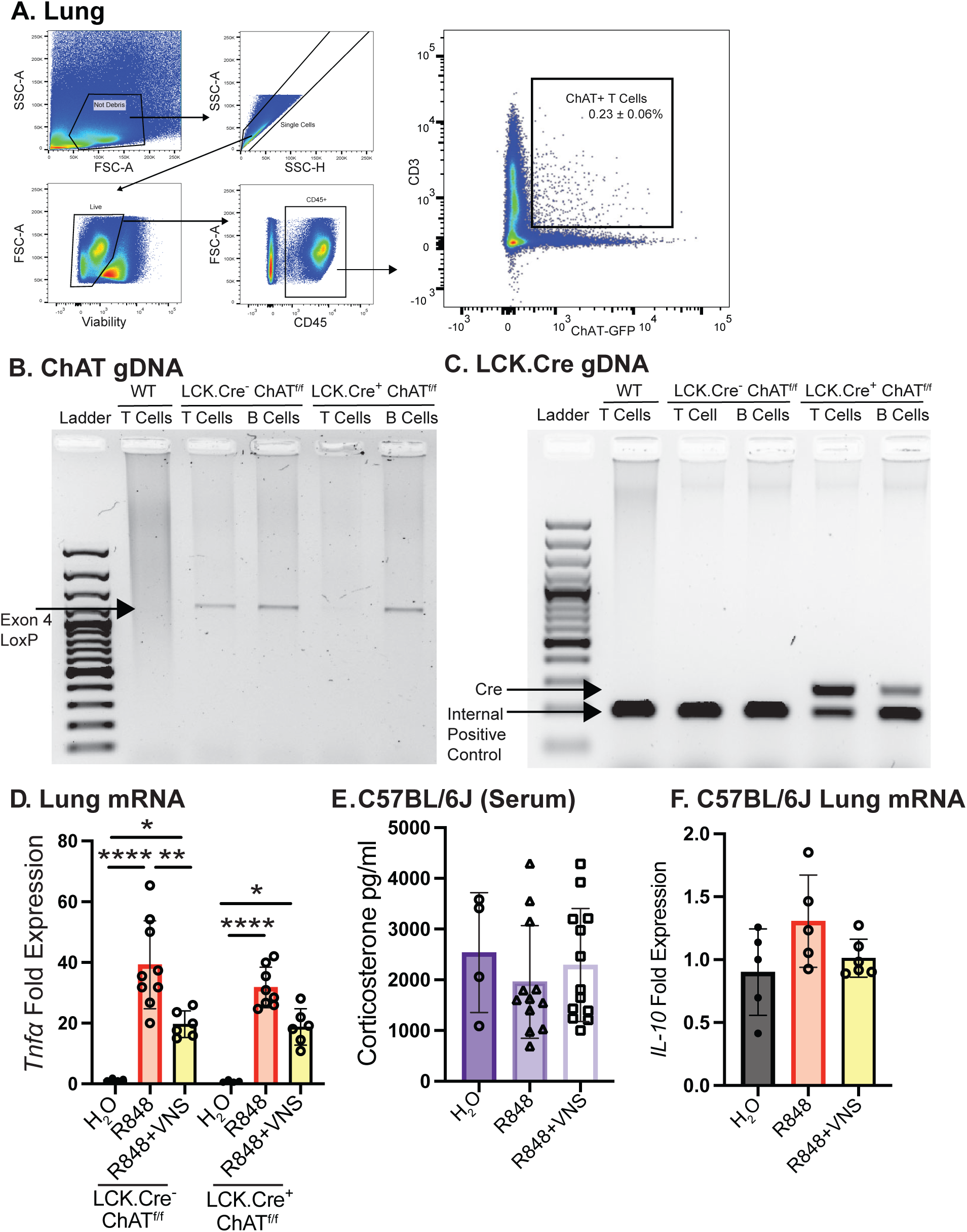
**(A)** Whole lungs were excised from ChAT-GFP mice and analyzed for frequency of ChAT+ T cells in the lung. **(B&C)** PCR on genomic DNA extracted from sorted splenic T and B cells. **(B)** Forward primer binds to exon 3 and reverse binds downstream of exon 4 in the ChAT gene with an expected product at 1.4kb or no product if sequence has been excised by Cre recombination. **(C)** Same DNA samples from **(B)** but run with primers used to identify Lck.Cre transgene. Expected product for Lck.Cre at 300bp and control band at 200bp. **(D)** Right lung cranial lobe *Tnfα* expression in LCK.Cre^-^ ChAT^F/F^ (WT) and LCK.Cre^+^ ChAT^F/F^ (T-cell cKO) mice subjected to VNS. **(E)** Serum corticosterone by commercial ELISA kit post VNS. **(F)** Right lung cranial lobe mRNA expression of *Il-10* post VNS. Data represented as mean ± standard deviation (SD). One-way ANOVA was used for statistical analysis followed by post-hoc analysis with Tukey’s multiple comparisons test. *p< 0.05; **p<0.01; ***p<0.001.

**Figure S3.**
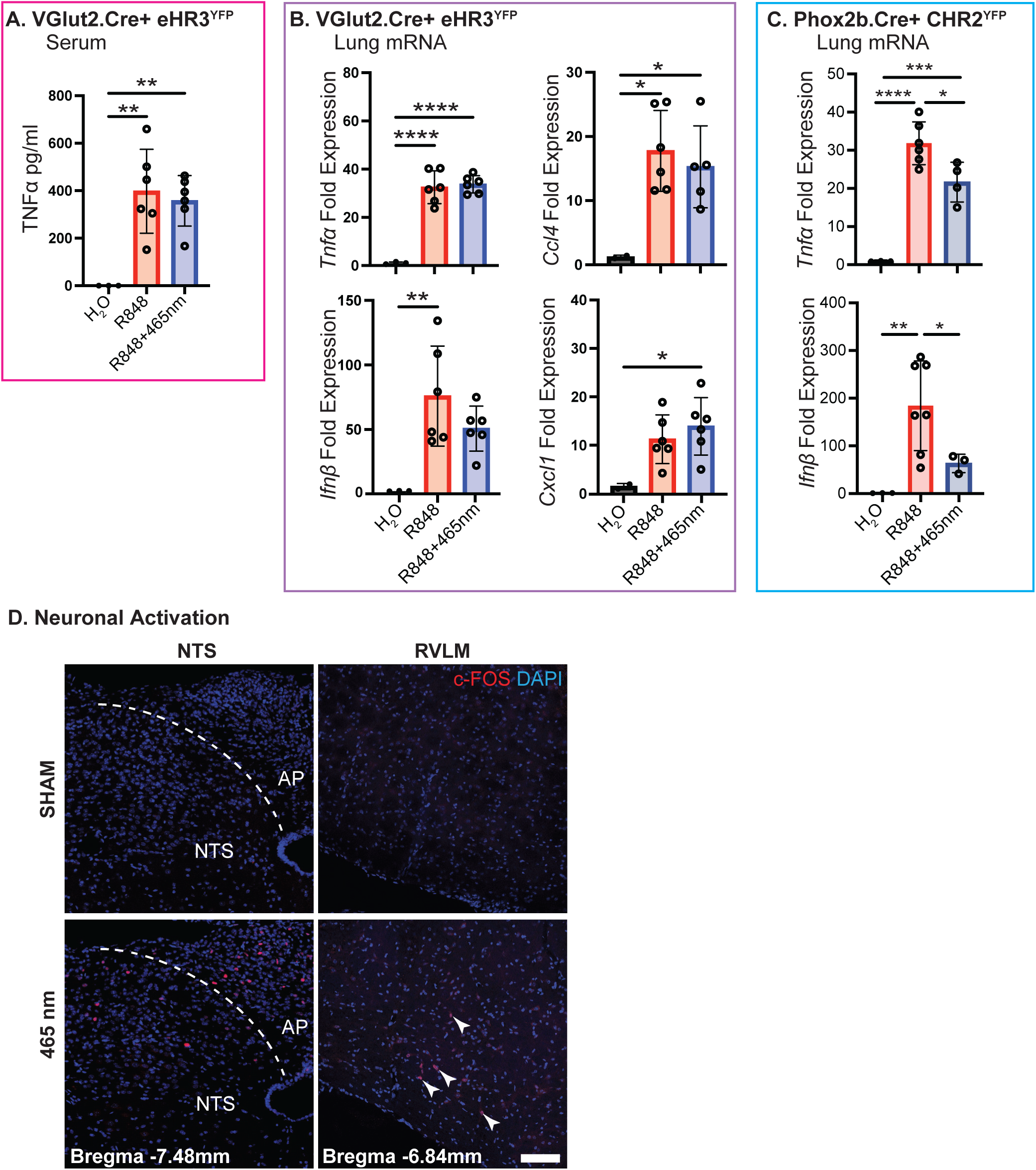
Optogenetic control experiments. The right nodose ganglion of VGlut2.Cre+ eHR3.YFP mice was exposed, and stimulated at 465 nm (inappropriate wavelength for eHR3) for 20 minutes. R848 was instilled 10 minutes into 465 nm stimulation, and mice were euthanized 1-hour post-R848 challenge. **(A)** TNFα protein was detected by ELISA and **(B)** lung mRNA expression of *Tnfα*, *Ifnp*, *Ccl4*, and *Cxcl1* was detected by qRT-PCR. Additional optogenetic stimulation of right nodose ganglia in wild type (C57BL/6J) vs. transgenic optogenetic (Phox2b.cre+ CHR2YFP mice) at 465 nm was then performed following the same experimental parameters. **(C)** Lung mRNA expression of Tnfα and Ifnβ by qRT-PCR **(D)** Activated (c-FOS+DAPI+) neurons in the NTS and RVLM of stimulated control C57BL/6J mice vs. Phox2b.Cre+ CHR2^YFP^ mice. Scale bar is 100 µm. Data represented as mean ± standard deviation (SD). One-way ANOVA was used for statistical analysis followed by post-hoc analysis with Tukey’s multiple comparisons test. Brown-Forsythe and Welch ANOVA was used for statistical analysis of Ifnβ in (C).*p< 0.05; **p<0.01; ***p<0.001.

**Figure S4.**
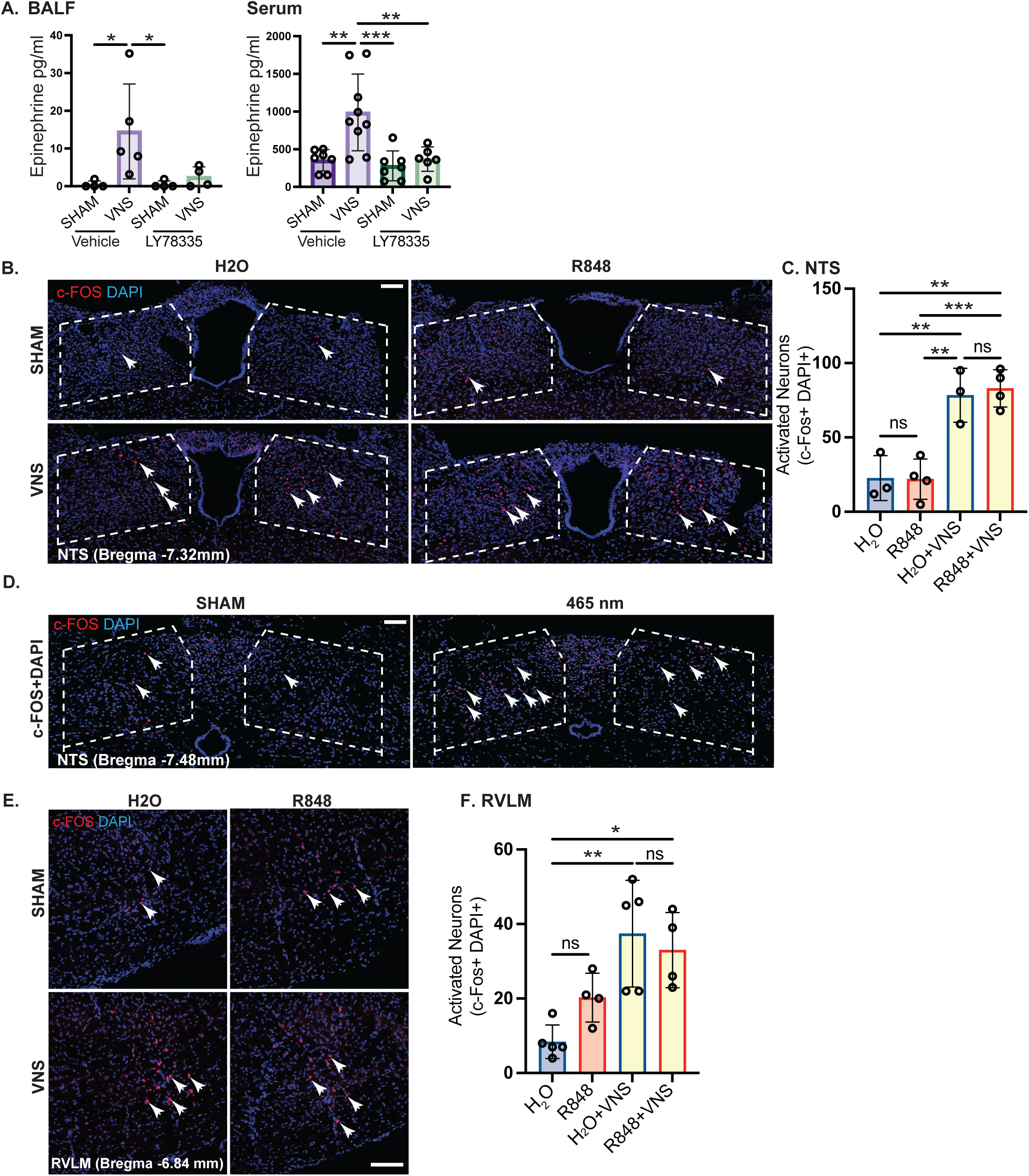
**(A)** PNMT inhibitor LY78335 or vehicle control (water) was i.p. injected 60 minutes prior to performing 5 minutes of VNS. Epinephrine was measured via commercial ELISA kit in serum and BALF. **(B)** Confocal image of NTS (positive regions outlined) in mice subject to 20 minutes of electrical VNS or SHAM stimulation. Scale bar is 100 µm. **(C)** Quantification of activated neurons within the NTS, determined by colocalization of c-FOS and DAPI. **(D)** Confocal image of NTS (positive region outlined) in mice subject to 465 nm optogenetic or control (590 nm) stimulation. Scale bar is 100 µm. **(E)** Confocal image and **(F)** quantification of right RVLM in mice subject to 20 minutes of electrical VNS or SHAM stimulation. Scale bar is 100 µm. Neuronal activation was determined by colocalization of c-FOS and DAPI. Data represented as mean ± standard deviation (SD). One-way ANOVA was used for statistical analysis followed by post-hoc analysis with Tukey’s multiple comparisons test. *p< 0.05; **p<0.01; ***p<0.001.

**Figure S5.**
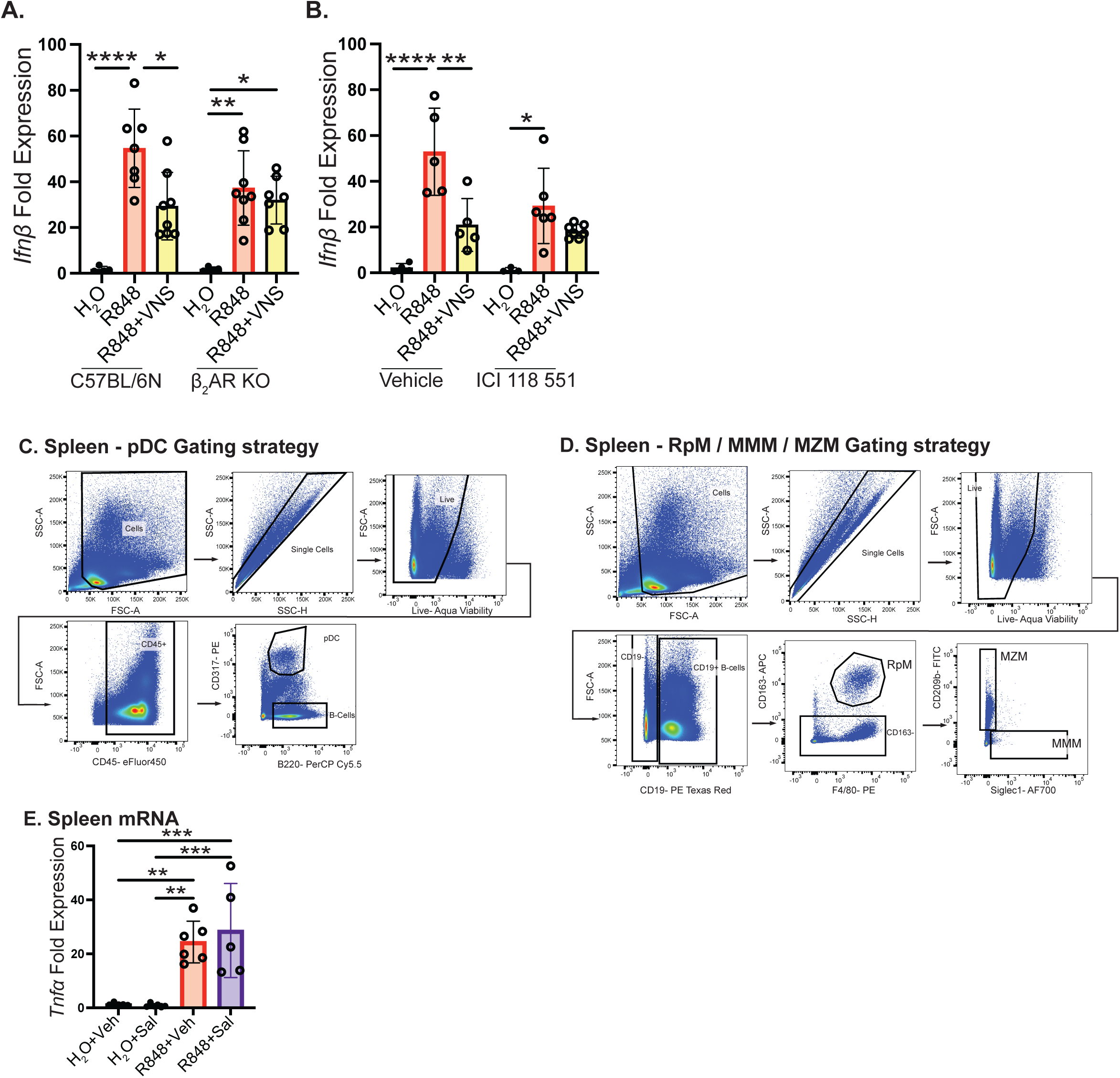
**(A)** Lung mRNA expression detected by qRT-PCR of *Ifnp* in wild-type (C57BL/6N) and β_2_AR knockout mice subjected to electrical VNS. **(B)** Lung mRNA expression of *Ifnp* in mice subjected to instillation of vehicle (water) or β_2_AR antagonist (ICI 118 551), prior to challenge with R848 and electrical VNS. **(C)** Gating strategy to distinguish pDC populations from spleen. **(D)** Gating strategy to distinguish, MMMφ, RPMφ, and MZMφ from the spleen. **(E)** Splenic mRNA expression of *Tnfα* in mice subject to R848 instillation with and without Salbutamol (1 mg/kg, i.v.). Data represented as mean ± standard deviation (SD). One-way ANOVA was used for statistical analysis followed by post-hoc analysis with Tukey’s multiple comparisons test. *p< 0.05; **p<0.01; ***p<0.001; ****p<0.0001.

